# Robust Homeostasis of Cellular Cholesterol via Antithetic Integral Control

**DOI:** 10.1101/2023.06.24.546224

**Authors:** Ronél Scheepers, Robyn P. Araujo

## Abstract

Although cholesterol is essential for cellular viability and proliferation, it is highly toxic in excess. The concentration of cellular cholesterol must therefore be maintained within tight tolerances, and is thought to be subject to a stringent form of homeostasis known as Robust Perfect Adaptation (RPA). While much is known about the cellular signalling interactions involved in cholesterol regulation, the specific chemical reaction network structures that might be responsible for the robust homeostatic regulation of cellular cholesterol have been entirely unclear until now. In particular, the molecular mechanisms responsible for sensing excess whole-cell cholesterol levels have not been identified previously, and no mathematical models to date have been able to capture an integral control implementation that could impose RPA on cellular cholesterol. Here we provide a detailed mathematical description of cholesterol regulation pathways in terms of biochemical reactions, based on an extensive review of experimental and clinical literature. We are able to decompose the associated chemical reaction network structures into several independent subnetworks, one of which is responsible for conferring RPA on several intracellular forms of cholesterol. Remarkably, our analysis reveals that RPA in the cholesterol concentration in the endoplasmic reticulum (ER) is almost certainly due to a well-characterised control strategy known as antithetic integral control which, in this case, involves the high-affinity binding of a multi-molecular transcription factor complex with cholesterol molecules that are excluded from the ER membrane. Our model provides a detailed framework for exploring the necessary biochemical conditions for robust homeostatic control of essential and tightly regulated cellular molecules such as cholesterol.

## 1 INTRODUCTION

Cholesterol is an amphipathic lipid that plays an essential role in regulating the properties of cell membranes in mammalian cells. Specifically, cholesterol forms part of the assembly and function of lipid micro– environments on the cell surface, known as lipid rafts (Simons and Toomre, 2000; Ouweneel et al., 2020). Enriched with free cholesterol (FC) and glycosphingolipids, these dynamic micro–domains function to segregate and distribute lipids and proteins to the cell surface. Here, the lipid rafts act as a platform for cellular receptors through which cellular processes such as endocytosis, exocytosis, receptor trafficking and cell signalling can be realised. (Simons and Ikonen, 2000; Simons and Toomre, 2000). Signalling pathways that involve lipid rafts include immunoglobulin E (IgE) signalling during the allergic immune response (Sheets et al., 1999; Simons and Toomre, 2000), T–cell antigen receptor signalling (Janes et al., 2000; Simons and Toomre, 2000) and Hedgehog signalling (Incardona and Eaton, 2000; Simons and Toomre, 2000).

In contrast, the altered composition of lipid rafts (Kulkarni et al., 2022) is characteristic of neurodegenerative diseases such as Parkinson’s, Alzheimer’s, multiple sclerosis and lysosomal storage disease (Kulkarni et al., 2022), while pathogens such as HIV, for example, exploit lipid raft composition to enter and exit its host cell (Simons and Ikonen, 2000). Glycosphingolipids and cholesterol are central players in lipid raft biology (Sonnino and Prinetti, 2013; Grassi et al., 2020), and considering the impacts of disrupted lipid raft composition, the total cellular level of cholesterol and its distribution between membranes, and within a given membrane, must be precisely controlled (Simons and Ikonen, 2000). Furthermore, as a macronutrient, cholesterol plays an integral role in the synthesis of steroid hormones and vitamin D, while dysregulated cholesterol metabolism can lead to pathophysiological diseases such age–related macular degeneration and cardiovascular disease (Scheepers et al., 2020; Curcio et al., 2011; Mc Auley and Mooney, 2014).

Evolution has endowed mammalian cells with intricate regulatory mechanisms to control cholesterol concentrations, ensuring adequate availability when required and shutting down cholesterol synthesis and uptake in conditions of cholesterol excess (Howe et al., 2016). The regulation of cholesterol takes place primarily in the plasma membrane (PM), endoplasmic reticulum (ER) and mitochondria, where the interplay between processes such as *de novo* biosynthesis, ingestion, efflux, storage and chemical modifications for specialised functions maintain steady cellular cholesterol levels (Tabas, 2002; Lange and Steck, 2016; Luo et al., 2020). Cholesterol regulation in the ER involves key proteins that respond to abundant cellular cholesterol levels by inhibiting cholesterol–promoting transcription pathways. Specifically, a family of transcription factors, collectively known as sterol regulatory element–binding proteins (SREBP), are sequestered by an ER–bound protein, Scap, to prevent gene transcription of the cholesterol biosynthesis pathway as well as the exogenous uptake of cholesterol from the circulation and homeostasis of cholesterol is restored. Post translationally, oxygenated derivatives of excess cholesterol activate the reverse cholesterol removal pathways to remove excess cholesterol and maintain stable cholesterol levels (Howe et al., 2016).

But despite the growing literature on the many metabolic and signal transduction events involved in cholesterol regulation, and although some mathematical models of cholesterol metabolism have been formulated (Paalvast et al., 2015; Pool et al., 2018), there is currently no comprehensive mathematical description of the molecular network mechanisms that can tightly control intracellular cholesterol concentration within sub–cellular compartments, and maintain these concentrations within very tight concentration tolerances, in the face of significant variations in the delivery and intracellular uptake of cholesterol.

The homeostasis imposed on subcellular cholesterol concentrations corresponds to a stringent form of regulation known as Robust Perfect Adaptation (RPA) (Araujo and Liotta, 2018, 2023b; Khammash, 2021). RPA is a biological network property whereby one or more molecular concentrations within the network are maintained at a particular fixed ‘setpoint’ at steady state, independently of specific network parameter choices (i.e., robustly), and in spite of altered inputs or disturbances to the system (Cannon, 1929; Kitano, 2007; Aoki et al., 2019; Ang et al., 2010; Ferrell, 2016; Jeynes-Smith and Araujo, 2021, 2023). The mathematical principles underpinning RPA in complex biological networks are now well understood, and we now have access to a universal description of RPA at both the macroscale of biochemical reaction networks (Araujo and Liotta, 2018, 2023a), and at the network microscale of individual intermolecular interactions and chemical reactions (Araujo and Liotta, 2023b). In particular, it is now known that all RPA-capable networks, no matter how large or complex, are decomposable into two distinct and well-defined subnetwork modules - Opposer modules and Balancer modules (Araujo and Liotta, 2018, 2023a). Whereas Opposer modules have an overarching feedback structure, into which special types of interactions (known as ‘opposer nodes’) are embedded, Balancer modules comprise a number of incoherent parallel pathways, incorporating specialised classes of interactions known as ‘balancer nodes’ and ‘connector nodes’. Particular network realisations of these two modular types can potentially be highly complex, and large and highly complicated RPA-capable networks can be constructed from the interconnections of these special RPA-conferring modular building blocks. We refer interested readers to (Araujo and Liotta, 2018), (Araujo and Liotta, 2023a), and (Khammash, 2021) for a comprehensive overview of the macroscale topological properties of RPA-capable networks.

The microscale RPA problem, which considers the detailed chemical reaction networks (CRNs) that can implement RPA through potentially intricate intermolecular interactions, has also now been solved (Araujo and Liotta, 2023b). In particular, it is now apparent that networks of chemical reactions that can support RPA are subject to very stringent structural criteria, which allow topological features (opposer nodes, balancer nodes, connector nodes) of an associated RPA-module to be created by linear integral controllers. In this way, RPA-conferring intermolecular interactions at the network microscale can be decomposed into a collection of subsidiary RPA-problems, each obtained by a linear coordinate change, that collaborate to impose RPA on the concentrations of certain specific molecules in the CRN.

In the present study, we identify the molecular mechanisms responsible for conferring RPA on plasma membrane cholesterol. In particular, we identify (i) the topological characteristics of the RPA-conferring network responsible for cholesterol homeostasis, (ii) the molecular mechanism by which cells sense excess cell cholesterol levels, and (iii) the integral that confers RPA on membrane-bound cellular cholesterol. In Section 2, we begin with an extensive review of the known molecular interactions and reactions involved in cholesterol regulation, including the critical transcriptional, translational and post-translational signalling events that regulate cholesterol within and between different subcellular compartments and organelles. In Section 3, we encode these detailed molecular mechanisms into a chemical reaction network (CRN), whose graphical structure and modularity can be analysed, and which can be used to identify molecular mechanisms directly responsible for RPA (Section 4).

## 2 THE CELL BIOLOGY OF CHOLESTEROL

### 2.1 Cholesterol regulation pathways

#### 2.1.1 Cholesterol flux

The ER contains an elaborate feedback system that is sensitive to the level of its membrane cholesterol by virtue of its small pool of cholesterol, which is set by the balance of PM cholesterol fluxes moving rapidly to and from the ER (Radhakrishnan et al., 2008; Lange and Steck, 2016). While little cholesterol moves from the plasma membrane to the ER at homeostatic cholesterol levels, the flux increases when plasma membrane cholesterol levels rise to the point where active cholesterol molecules, those above the association capacity of the bilayer phospholipids, escape from the surface of the bilayer and redistributes to the ER (Lange and Steck, 2020).

A proportion of cholesterol in the ER is converted to cholesterol esters (CE) by the ER enzyme acyl-coenzyme A (CoA):cholesterol acyltransferase (ACAT) (Chang et al., 2006, 2009). Newly synthesised CE accumulates within the rough ER and buds off as cytoplasmic lipid droplets, whereby it acts as a reservoir from which its hydrolysis can mobilise unesterified cholesterol for steroid hormone, oxysterols or bile acid production (Scheepers et al., 2020; Chang et al., 2009; Simons and Ikonen, 2000).

Reverse cholesterol transport begins with the formation of small high–density lipoprotein (HDL) particles by the liver and intestine. HDL acts as unesterified cholesterol acceptor that transports excess cholesterol within peripheral tissues to the plasma, a process mediated by ABCA1. Mature HDL particles then transport the cholesterol to the liver where it can be directly excreted into the bile or be metabolised into bile acids/salts before excretion (Feingold and Grunfeld, 2000; Ouimet et al., 2019).

Intracellular cholesterol is provided to the cellular organelles via biosynthesis and the uptake of circulating lipoprotein complexes (LDL) via lipoprotein receptors (LDLR) in the plasma membrane. While the balance between these two pathways depends on cell type and the availability of LDL–derived cholesterol, the master regulator of gene expression to realise these pathways is a polytopic ER membrane protein called Scap (Brown et al., 2018).

#### 2.1.2 SREBP transcription factors

Scap is inherently bound to sterol regulatory element–binding proteins (SREBP), a family of transcription factors anchored in the ER. Under abundant cholesterol conditions, the complexed Scap molecule binds to cholesterol and the hydrophobic ER protein Insig–1 (Brown et al., 2018; Gong et al., 2006). The conformational changes that Scap undergoes during this binding process prevent incorporation of the complex into COPII–coated vesicles for transport to the Golgi for processing (Sun et al., 2007). Consequently, the activation of gene transcription pathways further downstream is inhibited, biosynthesis and exogenous cholesterol uptake repressed, and cholesterol homeostasis is restored (Brown et al., 2018; Howe et al., 2016; Gong et al., 2006).

While *in vitro* studies confirmed the inhibition of SREBP processing under abundant cholesterol conditions, the mechanism by which the cell senses this abundance in the ER *in vivo* is not known (Lange and Steck, 1997). Motamed *et al*.(Motamed et al., 2011) showed the cholesterol-binding site in Scap is located in a membrane–associated loop that projects into the lumen of ER. Recognising that the study of Scap’s interactions with membrane cholesterol is technically difficult (Radhakrishnan et al., 2004), Gay *et al*. (Gay et al., 2015) used another sensor molecule, the soluble bacterial toxin perfringolysin O (PFO), to confirm the inhibition of SREBP processing and gene expression as reported by (Radhakrishnan et al., 2008). Based on this evidence, it was hypothesised that Scap may be binding to a pool of ER cholesterol that exceeds the sequestration capacity of the bilayer phospholipids and projects into the ER lumen, the exact location where the cholesterol–sensing domain of Scap resides (Motamed et al., 2011; Gay et al., 2015).

Under cholesterol–depleted conditions, Insig–1 dissociates from the SREBP/Scap complex to be ubiq-uitylated and degraded (Gong et al., 2006). The Scap/SREBP complex is now free to be carried to the Golgi complex, where fusion with the Golgi membrane results in proteolysis of SREBPs to release its transcription factors (TF). This nuclear fragment is translocated to the nucleus to, amongst many other activities, activate genes for the expression of (i) enzyme 3–hydroxy–3–methylglutaryl–coenzyme A reductase (HMGCR), (ii) low–density lipoprotein receptor (LDLR) on the plasma membrane, and (iii) proprotein convertase subtilisin/kexin type–9 (PCSK9) (Howe et al., 2016; Brown et al., 2018; Lagace, 2014).

#### 2.1.3 Cholesterol biosynthesis and down–regulation

The cholesterol biosynthesis process involves multiple complex pathways tightly controlled at various points, with the HMGCR enzyme catalysing the synthesis of mevalonate from 3–hydroxy–3–methylglutaryl– coenzyme A (HMG–CoA) in the first committed and rate–limiting step. This reaction forms the point of feedback control for the mevalonate pathway, and, as such, this enzyme is the primary target for medical intervention. Following a succession of intermediate changes, mevalonate is converted to lanosterol, the latter being converted to cholesterol after several sequential reactions. For the interested reader, Cerqueira *et al*. (Cerqueira et al., 2016) provides a detailed review of the biosynthesis of cholesterol.

The transcriptional down–regulation of cholesterol via the SREBP pathway is relatively slow, with the mRNA of target genes decreasing only after several hours. In contrast, 25–hydroxycholesterol (25HC) is an oxysterol produced by enzymatic conversion of ER cholesterol and is sensed by the membrane domain of HMG–CoA reductase to trigger its binding to Insig. Within minutes, this binding activates the enzyme’s ubiquitylation, degradation, and subsequent inhibition of cholesterol biosynthesis (Sharpe and Brown, 2013; DeBose-Boyd, 2008; Odnoshivkina et al., 2022).

#### 2.1.4 Receptor–mediated LDL acquisition

LDL receptors (LDLR) on the plasma membrane facilitate endocytosis of low–density lipoprotein (LDL) from the circulation. Endosomes release their content to lysosomes, where degradation of internalised LDL leads to the release of unesterified cholesterol and fatty acids (Brown and Goldstein, 1986). Recent studies by Trinh *et al*. (Trinh et al., 2020) confirmed that LDL–derived cholesterol from lysosomes are transported to the PM first to maintain optimal cholesterol levels, and subsequently from PM to the regulatory domain of the ER.

Transcriptional down–regulation of LDLR is facilitated by PCSK9, a soluble member of the proprotein convertase family of secretory serine endoproteases. Upon binding to LDLR on the cell surface, the complex is redirected for internalisation into lysosomes, where PCSK9 and LDLR are degraded. Consequently, LDLR recycling to the cell surface is prevented and LDL uptake from the circulation down–regulated (Lagace, 2014).

As an essential structural component of cell membranes, cholesterol modulates the properties of the membrane lipid bilayers by intercalating with phospholipids to maintain membrane fluidity, and rigidity (Lange and Steck, 2020). The association of cholesterol with diverse organelle phospholipids in specific ratios in cell membranes has been described by McConnell and Radhakrishnan (McConnell and Radhakrishnan, 2003) as *stoichiometric complexes*. Association equilibrium is achieved when cholesterol accumulates as phospholipid complexes up to a particular stoichiometry or set–point (Lange et al., 2013; Yeagle and Young, 1986). Conversely, uncomplexed sterol molecules, those above the association capacity of the bilayer phospholipids, have a relatively high tendency to escape from the surface of the bilayer. This tendency is also referred to as a constituent’s *activity* or *fugacity* (Lange and Steck, 2008) and this fraction of cholesterol is called *active cholesterol* (Lange and Steck, 2016). These *active* molecules rapidly redistribute to the cytoplasmic membranes down its activity gradient via Aster proteins (Sandhu et al., 2018), where a rise in intracellular pools adjusts the synthesis and uptake of cholesterol to return the plasma membrane cholesterol to its physiological set–point (Lange and Steck, 2008, 2016).

#### 2.1.5 Liver X (LXR) transcription factors

27–hydroxycholesterol (27HC), another oxysterol derivative of ER cholesterol, is a transcriptional activator of a member of the nuclear receptor superfamily, the liver X receptor (LXR). This subfamily of transcription factors consists of two distinct members, LXR*β*, being ubiquitously expressed, and LXR*α*, whose expression is restricted to tissues rich in lipid metabolism (Lu et al., 2001). Being a Type–II receptor, LXR resides in the nucleus and is bound to its specific DNA response elements, even in the absence of ligand.

27HC mediates signal transduction through direct binding to LXR in the nucleus at physiological concentrations (Howe et al., 2016; Janowski et al., 1999; Sever and Glass, 2013). Once activated, LXR upregulates ATP–binding cassette transporter genes (ABCA1) to facilitate cholesterol efflux via systemic high–density lipoprotein (HDL) particles. Furthermore, activated LXR upregulates the inducible degrader of LDLR (Idol), to decrease cholesterol uptake from LDL (Howe et al., 2016; Zelcer et al., 2009).

## 3 METHODS

### 3.1 Chemical reaction network representation of cellular cholesterol regulation

Here we represent the biochemical processes reviewed in detail in Section 2 as a collection of chemical reactions, which gives rise to a CRN involving fourteen molecular species, comprising transcription factors, proteins, biochemical intermediates and macro–molecules (see Table 1). These key molecules participate in twenty-two chemical reactions (see Table 2). One of the species — the (extracellular) concentration of cholesterol–carrying lipoproteins (*C*_*L*_) — serves as the distinguished input molecule for this system, since this determines the delivery of cholesterol to the cell, and is therefore the ‘stimulus’ to which intracellular cholesterol must adapt and exhibit homeostasis/RPA.

**Table 1.** List of species and associated symbols

| Species | Symbol |
| --- | --- |
| Active cholesterol | $C$ |
| Cholesterol in ER membrane | $C_e$ |
| Cholesterol in PM | $C_p$ |
| Cholesterol-carrying LDL | $C_L$ |
| LDL receptor (LDLR) | $R$ |
| Cholesterol released from LDLR | $C_f$ |
| Esterified cholesterol | $E$ |
| 3-hydroxy-3-methylglutaryl-coenzyme A (HMG-CoA) | $H$ |
| 3-hydroxy-3-methylglutaryl-coenzyme A reductase (HMGCR) | $H_R$ |
| proprotein convertase subtilisin/kexin type-9 (PCSK9) | $P$ |
| SREBP transcription factor for LDLR | $S_r$ |
| SREBP transcription factor for HMGCR | $S_h$ |
| SREBP transcription factor for PCSK9 | $S_p$ |
| SREBP/Scap/Insig complex | $S_{ci}$ |

**Table 2.**
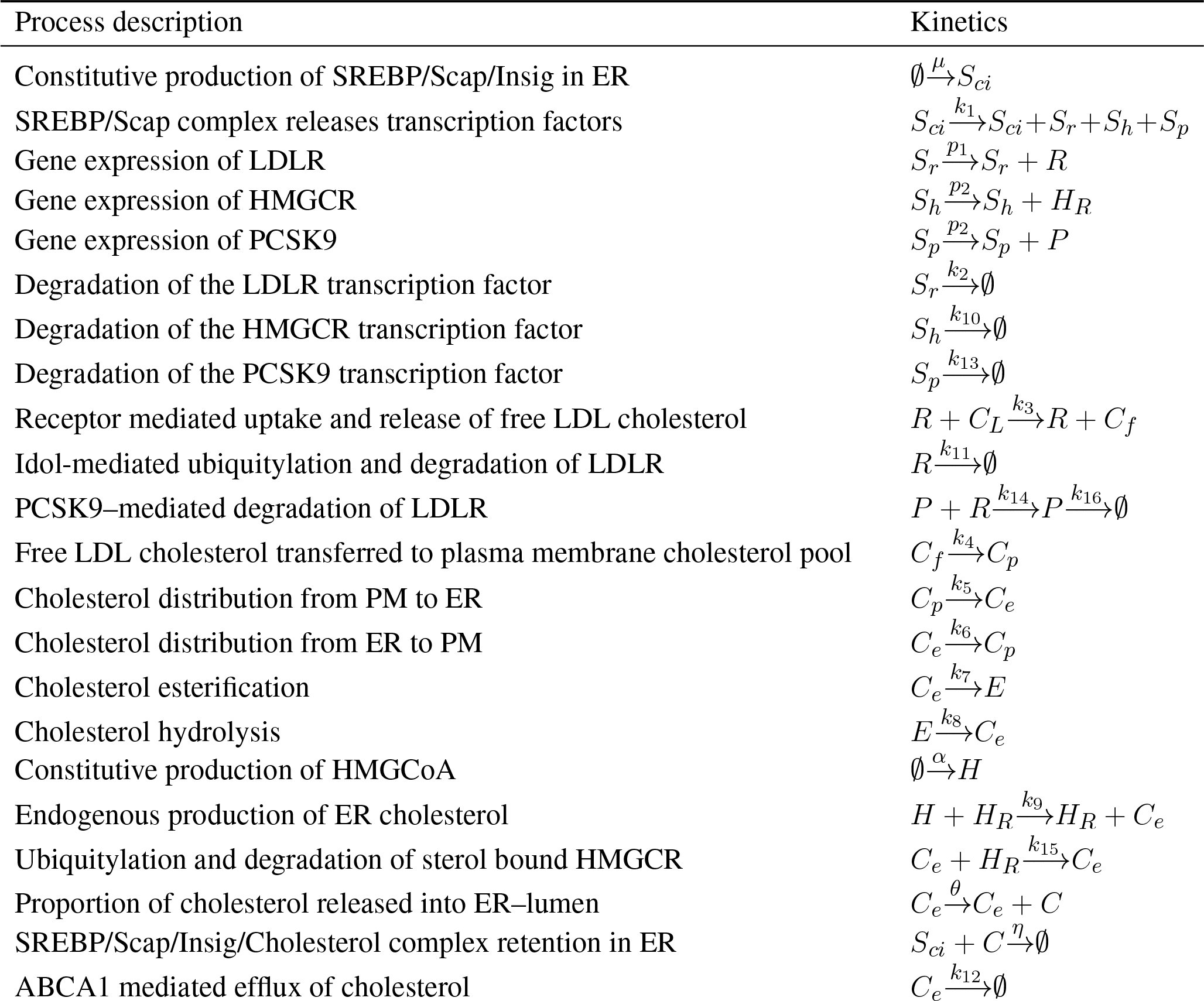
Listing of the individual reactions comprising the CRN for cellular cholesterol regulation.

The biochemical processes captured by the individual reactions of the CRN are as follows:

We capture the constitutive production rate of the SREBP/SCAP/Insig1 complex in the ER by the reaction 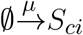.

Under cholesterol–depleted conditions, *S*_*ci*_ complexes are released to the Golgi for processing (dissociation and degradation of Insig–1 are omitted from our model). Proteolytic activity in the Golgi ensures the release of the SREBP transcription factors *S*_*r*_, *S*_*h*_ and *S*_*p*_ to the nucleus, expressed here as 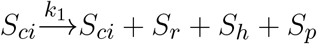.

The rate at which the transcription factor *S*_*r*_ activates transcription of its target genes to produce LDLR protein is captured as 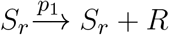. Similarly, *S*_*h*_ induces the production of HMGCR, 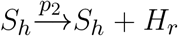, while the production of PCSK9 from *S*_*p*_ transcription is represented as 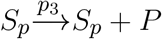. Degradation rates of the SREBP transcription factors are expressed as 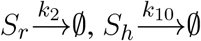 and 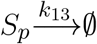 respectively (Sundqvist and Ericsson, 2003).

Receptor–mediated uptake and release of unesterified cholesterol via endosomes are captured in the reaction 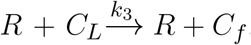. Here, the conservation of *R* represents the recycling of LDLR to the surface membrane. The interested reader is referred to (Tindall et al., 2009) and (Pool et al., 2018) for comprehensive modelling of this recycling process. Post–translational regulation of LDLR occurs via IDOL mediated degradation, simply captured as 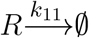, while the binding of PCSK9 to LDLR facilitates the degradation of LDLR as well as its own degradation in the lysosomes. These degradation rates are expressed by the equations 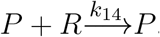, and 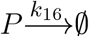, respectively.

*C*_*f*_ is transported to the PM, captured by the rate equation 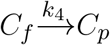. From here, fluxes between cholesterol in the PM, *C*_*p*_, and the ER membrane cholesterol pool (*C*_*e*_), are captured as 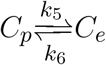.

A proportion of ER cholesterol is converted to cholesterol esters at a rate, 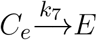, while a proportion of *E* can be converted back to unesterified ER cholesterol when required at a rate 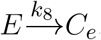.

Returning to the cholesterol biosynthesis pathway, the rate of constitutive production of HMGCoA (*H*) is expressed as 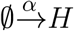.

Following a series of omitted intermediate steps, the final product in the biosynthesis pathway, *C*_*e*_, is produced via activation of HMGCoA by the HMGCR reductase enzyme, modelled as 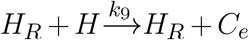.

While cholesterol biosynthesis is tightly regulated via various feedback processes that lead to the degradation of HMGCR, we represent this cholesterol initiated degradation process as 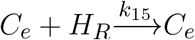.

ER cholesterol molecules exceeding the sequestration capacity of the phospholipid bilayer, *C*, is released into the ER lumen, modelled here by the rate equation 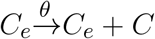.

The cholesterol molecules in the ER lumen are now sensed and bound by the *S*_*ci*_ complex, resulting in the irreversible retention of a *S*_*ci*_ *C* complex in the ER membrane, modelled as 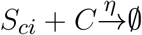. In this case, transport of the *S*_*ci*_ complex to the Golgi for release of SREBP transcription factors is inhibited, resulting in a decrease in cellular cholesterol concentration.

Finally, ER cholesterol efflux via systemic high–density lipoprotein (HDL) particles is represented as 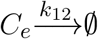.

A graph may be obtained from a set of chemical reactions by interpreting the multisets of species constituting either reactants or products (known collectively as ‘complexes’ in chemical reaction network theory (CRNT)) as vertices, and the reactions themselves as directed edges. Connected components of the graph are referred to as ‘linkages classes’ in CRNT (Feinberg, 2019). A ‘strong linkage class’ corresponds to a strongly connected component (SCC) of the graph, being a maximal strongly-connected subgraph. A ‘terminal strong linkage class’ is a strong linkage class in which no complex reacts to a complex in a different strong linkage class. Complexes belonging to terminal strong linkage classes are referred to as *terminal complexes*; all other complexes are *non-terminal complexes*.

For any such CRN graph structure, a key integer invariant known as the *deficiency* of the CRN may be computed, providing a quantitative measure of the linear independence of the CRN reactions given their distribution into linkage classes (Feinberg, 2019; Araujo and Liotta, 2023b).

The deficiency, *δ*, of a CRN is given by

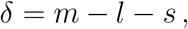

where *m* is the number of complexes, *l* is the number of linkage classes, and *s* is the dimension of the stoichiometric subspace (the span of the reaction vectors), also known as the rank of the CRN. In Figure 1 we organise the twenty-two reactions of the cholesterol homeostasis CRN into linkage classes. For this network, *m* = 23, *l* = 2 and *s* = 15, which gives a deficiency *δ* = 6.

**Figure 1.**
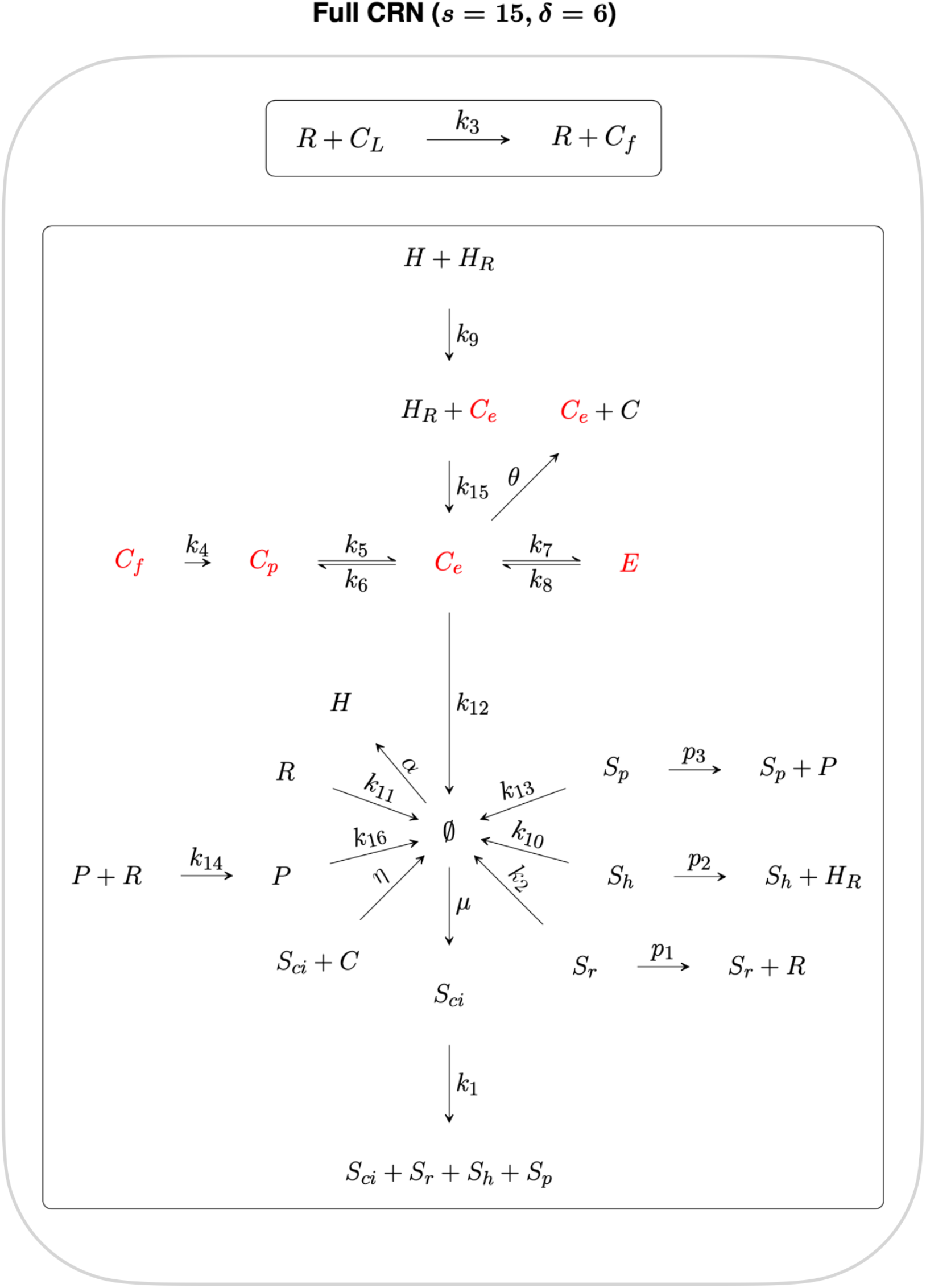
CRN graph structure for the cholesterol homeostasis network.

## 4 RESULTS

### 4.1 Graph analysis of the CRN

By decomposing this CRN graph into algebraically independent subnetworks, the CRN reactions can be partitioned into subsets whose steady states may be determined independently from the rest of the CRN.

In the context of RPA, such a decomposition into independent subnetworks allows us to distinguish the network reactions that contribute to the network’s RPA capacity from reactions that play no role in the RPA capacity of the CRN (Araujo and Liotta, 2023b).

Subnetworks are algebraically independent when their individual ranks sum to the rank of the parent network (Feinberg, 2019). Here we decompose the full CRN into two algebraically independent subnetworks, as shown in Figure 2 and Figure 3, where the sum of the ranks of the two subnetworks (*s*_1_ = 5 and *s*_2_ = 10 respectively) gives the rank of the full CRN (*s* = 15). Furthermore, the sum of the respective subnetwork deficiencies, *δ*_1_ = 1 and *δ*_2_ = 5 gives the full CRN deficiency of *δ* = 6.

**Figure 2.**
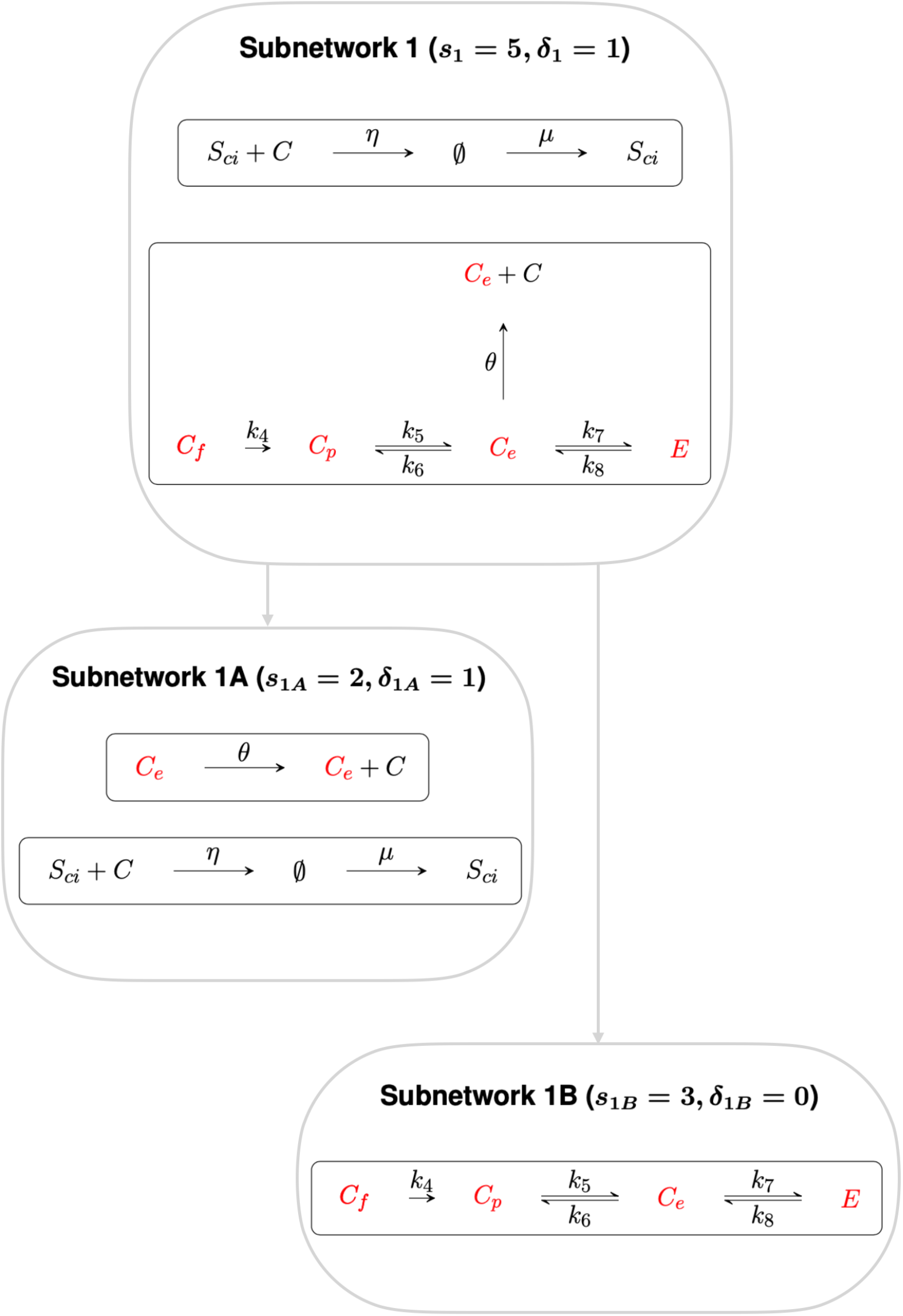
Subnetwork 1. Subnetwork 1, with deficiency 1, confers RPA on the species *C*_*e*_, *C*_*p*_, *C*_*f*_ and *E*. Subnetwork 1 can be decomposed into algebraically independent Subnetworks 1A and 1B respectively. Subnetwork 1A contains species *C*_*e*_, that regulates the transcription of cholesterol promoting processes, while Subnetwork 1B comprise the other three non–terminal complexes, *C*_*p*_, *C*_*f*_ and *E*.

**Figure 3.**
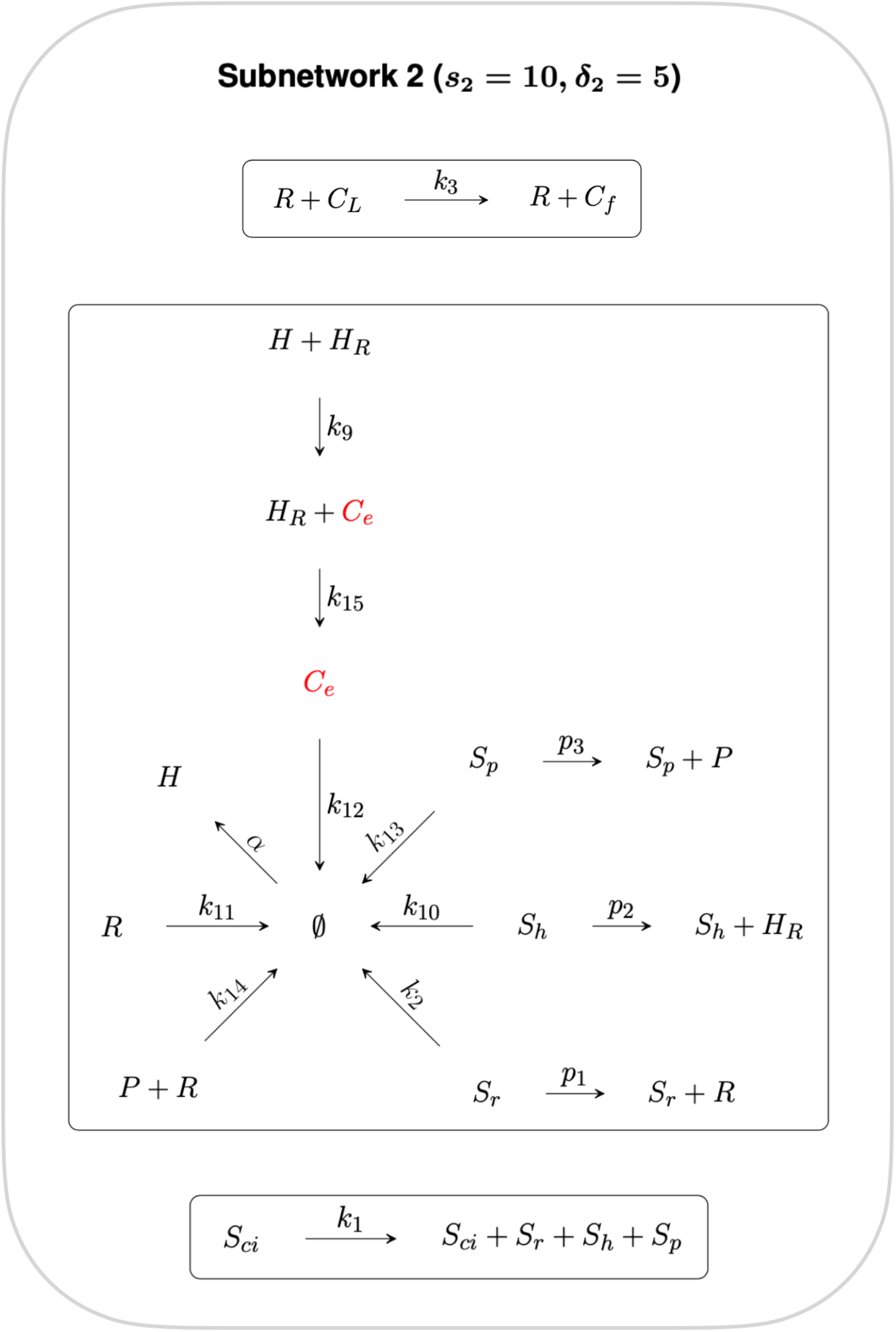
Subnetwork 2. Reactions in this Subnetwork contribute to the overarching controlling module but do not have any influence on the RPA capacity of the CRN as a whole. *C*_*e*_ is the only species common to both Independent Subnetworks, and thus connects independent CRN subnetworks 1 and 2.

Subnetwork 1 contains six non–terminal complexes *C*_*e*_, *C*_*p*_, *C*_*f*_, *E, S*_*ci*_ and ∅. CRNs with a deficiency of precisely one can, by the Shinar–Feinberg Theorem (Shinar and Feinberg, 2010), achieve RPA if the CRN contains two distinct non–terminal complexes that differ in a single species. Since ∅ is a non-terminal complex in this algebraically-independent deficiency-one subnetwork, all non-terminal complexes comprising a single species (*C*_*e*_, *C*_*p*_, *C*_*f*_ and *E*) are guaranteed to be RPA-capable.

We further decompose Subnetwork 1 into Subnetworks 1A and 1B, where the deficiency of Subnetwork 1A and Subnetwork 1B is *δ*_1*A*_ = 1 and *δ*_1*B*_ = 0 respectively. Subnetwork 1A contains the species involved in regulating the transcription of cholesterol–promoting processes via ‘entrapment’ of the *S*_*ci*_*C* complex in the ER, including the non-terminal complex *C*_*e*_. Applying the Shinar–Feinberg Theorem (Shinar and Feinberg, 2010) to Subnetwork 1A, the two non–terminal complexes *C*_*e*_ and ∅ differ in the single species *C*_*e*_ only, identifying *C*_*e*_ as the species capable of conferring RPA to the cellular cholesterol network. Subnetwork 1B contains the other three non–terminal complexes, *C*_*p*_, *C*_*f*_ and *E*.

The reactions contained in Subnetwork 2 contribute to the overarching control of cellular cholesterol but do not have any influence on the RPA capacity of the CRN as a whole, and is therefore excluded from further analysis.

Having identified the controller reactions and the RPA–capable species of the cellular cholesterol network, we now proceed to determine the linear combination of rate equations that identify the integrator responsible for conferring RPA on the system, as well as the respective set points of each individual RPA species.

### 4.2 Mass action analysis

CRNs can also induce a set of rate equations under the mass–action assumption, whereby each reaction proceeds at a rate proportional to the concentration of each reactant. There now exist methods to analyse such polynomial dynamical systems to systematically extract the integral-control implementing features of RPA-capable CRNs (Araujo and Liotta, 2023b). For the cholesterol homeostasis CRN under consideration here, the mass-action rate equations are:

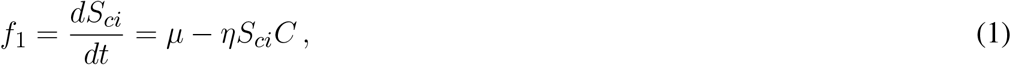

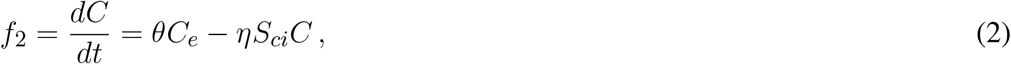

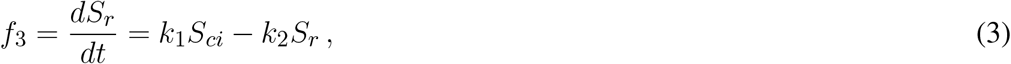

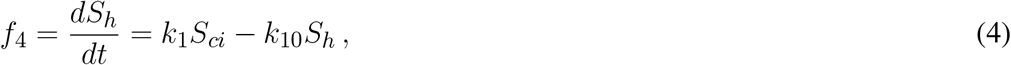

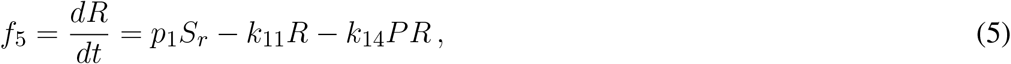

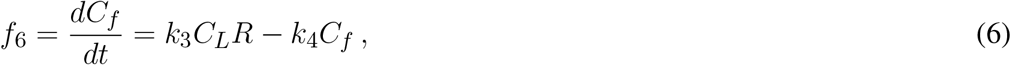

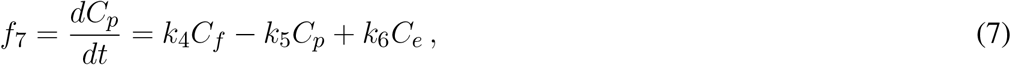

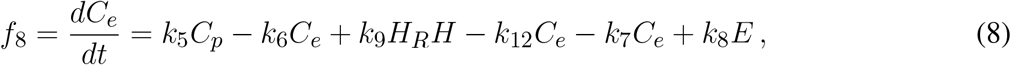

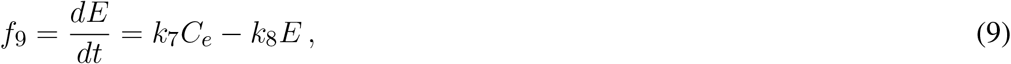

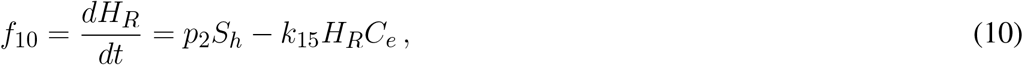

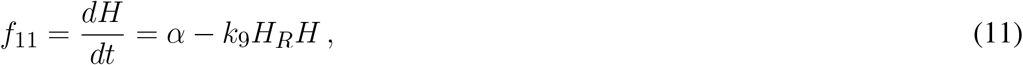

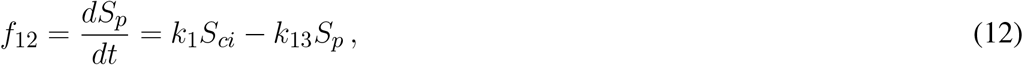

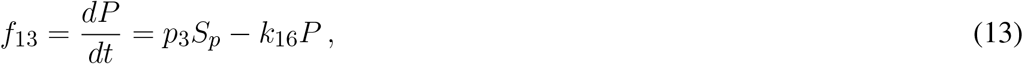

Theorem 1 in (Araujo and Liotta, 2023b) makes clear that for any RPA–capable CRN with interacting molecules *x*_1_, …*x*_*n*_ and corresponding mass–action rate equations *f*_1_, …*f*_*n*_, there always exists a linear combination of polynomials {*r*_1_, …, *r*_*n*_} ⊂ ℝ [*x*_1_, …, *x*_*n*_] such that

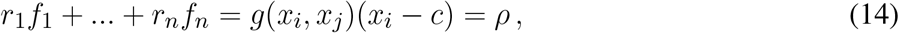

where *ρ* = *g*(*x*_*i*_, *x*_*j*_)(*x*_*i*_ − *c*), in its lowest form, is the RPA polynomial of the CRN, being a special polynomial function in two variables. When this occurs, the CRN has the capacity for RPA in the variable *x*_*i*_, with setpoint *x*_*i*_ = *c*, while *x*_*j*_ is any variable that does not exhibit RPA. Moreover, if *x*_*i*_ = *c* is a stable steady-state of the CRN’s mass-action equations, the CRN *exhibits* RPA in the variable *x*_*i*_. Theorem 1 ((Araujo and Liotta, 2023b)) makes a distinction between the variables and the species of a CRN kinetic model in cases where the rate equations contain a special type of monomial known as a *boundary variable* (see Supplementary Information S1.5 in (Araujo and Liotta, 2023b) for a full technical discussion); the rate equations under consideration here contain no boundary variables, however, so the variables of the model are the fourteen species. Thus, we seek a projection of the ideal ⟨*f*_1_, …, *f*_13_⟩ onto two species. We expect *C*_*e*_ to be RPA–capable for this model, based on the analysis in the preceding section. Moreover, the input to the system, *C*_*L*_, is not RPA–capable by definition. We therefore compute the elimination ideal ⟨*f*_1_, …, *f*_13_⟩ ∩ ℝ [*C*_*e*_, *C*_*L*_] via computation of a suitable Gröbner basis. For an efficient computation, we follow the method presented in (Araujo and Liotta, 2023b) which employs the block elimination order which partitions the species into two blocks — the two projection species (in this case {*C*_*e*_, *C*_*L*_}), and a second set containing all remaining species — with the *degree reverse lexicographic order* imposed on each block individually. We adapt the code provided in the Supplementary Information in (Araujo and Liotta, 2023b), which uses the open–source software *Singular* (https://www.singular.uni-kl.de/).

Using this method, we compute the two–variable elimination ideal to be

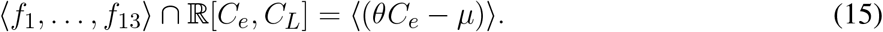

We note that the right–hand side of Equation 15 has the form of an RPA polynomial that is zero–order in the non–RPA-capable variable, *C*_*L*_, thereby confirming that the system is RPA–capable in the species *C*_*e*_, and also calculates that its steady–state setpoint is *C*_*e*_ = *μ/θ*. Setpoints for the variables *C*_*p*_, *C*_*f*_ and *E* can be computed similarly: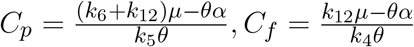 and 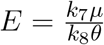, respectively. These calculations also confirm RPA capacity in all four variables suggested by the graphical analysis.

Moreover, the linear combination of rate equations used to identify the RPA polynomial (15) within the steady–state ideal can be obtained directly using the *lift* function in *Singular*. In this case, we compute that

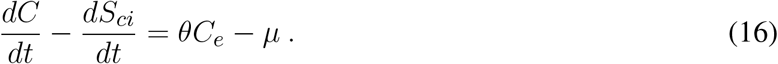

This calculation reveals the linear coordinate change that identifies an ‘internal model’ ((Araujo and Liotta, 2023a,b, 2018)), and demonstrates that the variable (*C* − *S*_*ci*_) is the integral variable (or ‘actuator’) for the integral controller. In addition, by computing the syzygies for the ideal, we confirm that this system has no linear syzygies. Thus, there are no mass conservation relationships embedded within the rate equations, and this linear coordinate change (16) is unique. We provide full details of these methods, along with an extended presentation of our analysis in our accompanying Supplementary Materials. Our *Singular* code is available at https://github.com/RonelScheepers/RobustPerfectAdaptation.

In Fig 4, we demonstrate via numerical simulations that the species *C*_*e*_, *C*_*f*_, *C*_*p*_ and *E* do indeed exhibit RPA for the indicated choice of parameters, and return to their expected setpoints for a range of step increases in the system input, *C*_*L*_. By contrast, none of the remaining nine species exhibits RPA, instead achieving a steady–state value that varies with *C*_*L*_. To confirm that these stable steady–state responses were achievable for a significant region of parameter space, we used Matlab’s symbolic toolbox to compute the characteristic polynomial for the system’s Jacobian matrix, and solved for the system’s eigenvalues using 10,000 different random parameter sets, with individual parameters selected from a uniform distribution on the interval (1, 50). This process was repeated for ten trials; for each trial, roughly 50% of parameter sets produced a stable system, with all calculated eigenvalues lying strictly in the left–half of the complex plane. This confirms that the CRN is not dependent on special fine–tuning of parameters to achieve stability, and can exhibit RPA in the noted variables for a wide range of different parameters. All Matlab code for this stability analysis is included at https://github.com/RonelScheepers/RobustPerfectAdaptation.

**Figure 4.**
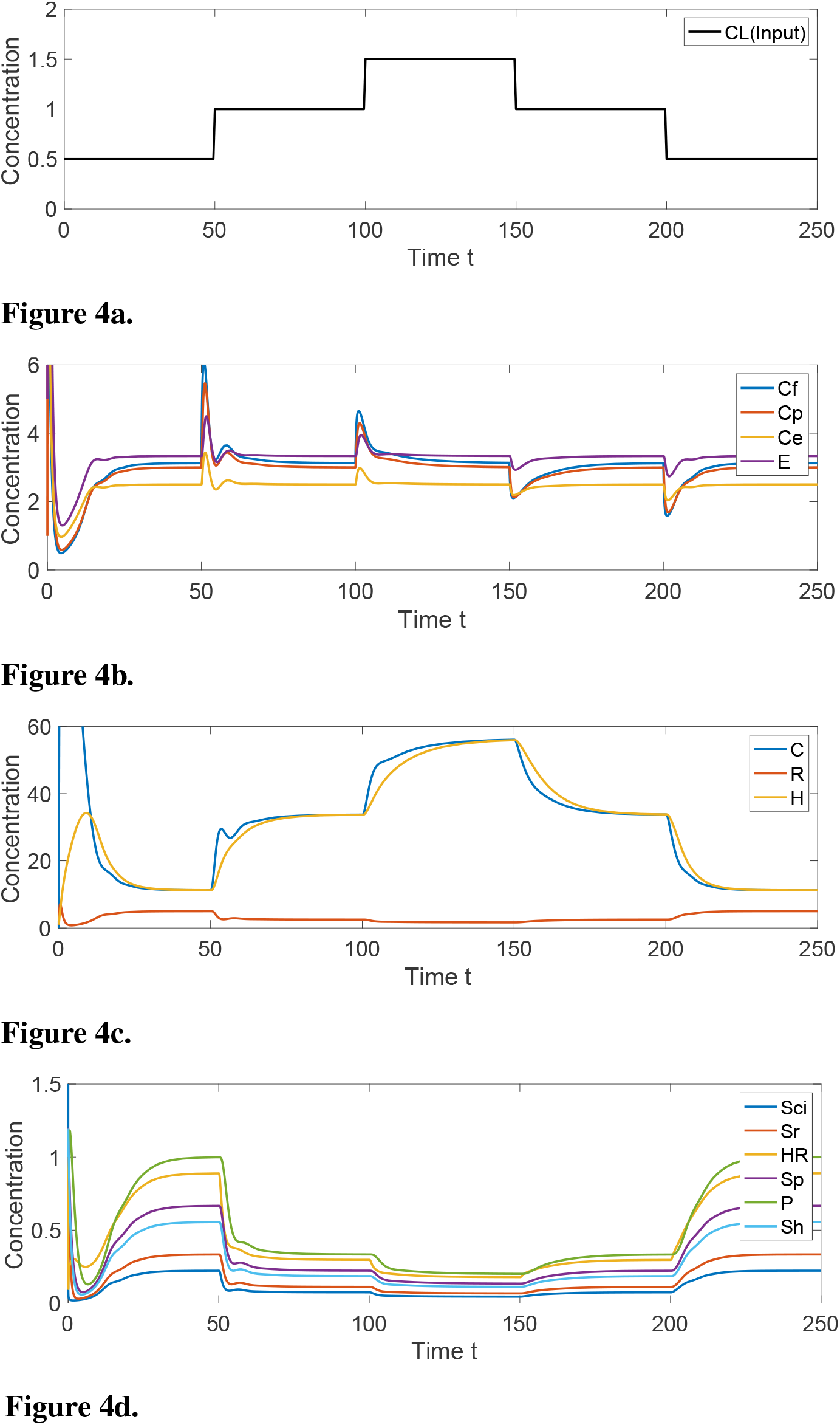
System response to step-changes in *C*_*L*_. **(4a)** Concentration of input variable, *C*_*L*_. **(4b)** For indicated persistent disturbances in the concentration of *C*_*L*_, the concentration of the RPA–capable variables *C*_*e*_, *C*_*f*_, *C*_*p*_ and *E* transiently change, and then return to their respective fixed setpoints. **(4c)** The non–RPA capable variables *C, R* and *H* arrive at steady-state values that depend on the value of *C*_*L*_. **(4d)** The non–RPA capable variables *S*_*ci*_, *S*_*r*_, *H*_*R*_, *S*_*p*_, *P* and *S*_*h*_ arrive at steady-state values that depend on the value of *C*_*L*_. Parameters: *k*_1_ = 3, *k*_2_ = 2, *k*_3_ = 5, *k*_4_ = 4, *k*_5_ = 5, *k*_6_ = 1, *k*_7_ = 4, *k*_8_ = 3, *k*_9_ = 1, *k*_10_ = 1.2, *k*_11_ = 1, *k*_12_ = 9, *k*_13_ = 1, *k*_14_ = 1, *k*_15_ = 1, *k*_16_ = 1, *p*_1_ = 30, *p*_2_ = 4, *p*_3_ = 1.5, *μ* = 25, *η* = 10, *α* = 10, *θ* = 10. Initial values: *S*_*ci*_(0) = 5, *C*(0) = 1, *S*_*r*_(0) = 1, *S*_*h*_(0) = 1, *R*(0) = 1, *C*_*f*_ (0) = 50, *C*_*p*_(0) = 1, *C*_*e*_(0) = 100, *E*(0) = 5, *H*_*R*_(0) = 1, *H*(0) = 1, *S*_*p*_(0) = 1, *P* (0) = 1.

## 5 DISCUSSION

The quest for the fundamental design principles that govern the robust implementation of important biological functions, particularly within the highly complex molecular interaction networks within single cells, is considered to be one of the most important grand challenges in the life sciences (Araujo and Liotta, 2023a,b). Robust Perfect Adaptation (RPA) is a keystone signalling phenomenon that is ubiquitously observed at all scales of biological organisation, and the only robust biological functionality for which there now exists a universal solution — at both the network macroscale (Araujo and Liotta, 2018) and the network microscale (Araujo and Liotta, 2023b). This universal solution makes clear that all RPA-capable networks are necessarily modular by design, and decomposable into subnetworks (‘modules’) from two and only two distinct classes — Opposer modules, with an overarching feedback structure; and Balancer modules, with an overarching feedforward structure. Moreover, these structures are characterised by well–defined topological features: opposer nodes (in Opposer modules), and balancer nodes and connector nodes (in Balancer modules). It is now known that, at the molecular level of biochemical reactions, all of these distinctive topological features (nodes) must be identifiable through linear coordinate changes in order for a chemical reaction network to exhibit RPA. In this way, control systems generated via intermolecular interactions and enzyme–catalysed reactions implement a special form of integral control that differs in important ways from modern engineering control systems that employ specially–designed components for the computation of integrals.

Although cellular cholesterol is thought to be subject to a very stringent form of homeostatic control, and thus RPA, the detailed molecular mechanisms responsible for such robust control have been unknown until now, owing largely to the relative complexity of the signalling network underpinning cholesterol regulation in comparison with most known RPA–capable chemical reaction networks (CRNs). In this study, we develop a CRN for cholesterol regulation that encompasses a comprehensive review of known molecular interactions involved in the regulation of intracellular cholesterol concentrations. This has allowed us to undertake a detailed analysis of the signalling architectures that are involved in the tight regulation of cholesterol concentrations within cellular and sub–cellular membranes, from both a graphical analysis of the CRN structure, as well as an algebro–geometric analysis of the corresponding mass–action equations.

Our analysis makes clear that the tight homeostatic control of membrane–bound cholesterol, in both the cellular plasma membrane and in the endoplasmic reticulum (ER), is the result of a single Opposer module, containing a single opposer node. We summarise the architecture of this network schematically in Figure 5, highlighting the overarching feedback architecture of the network.

**Figure 5.**
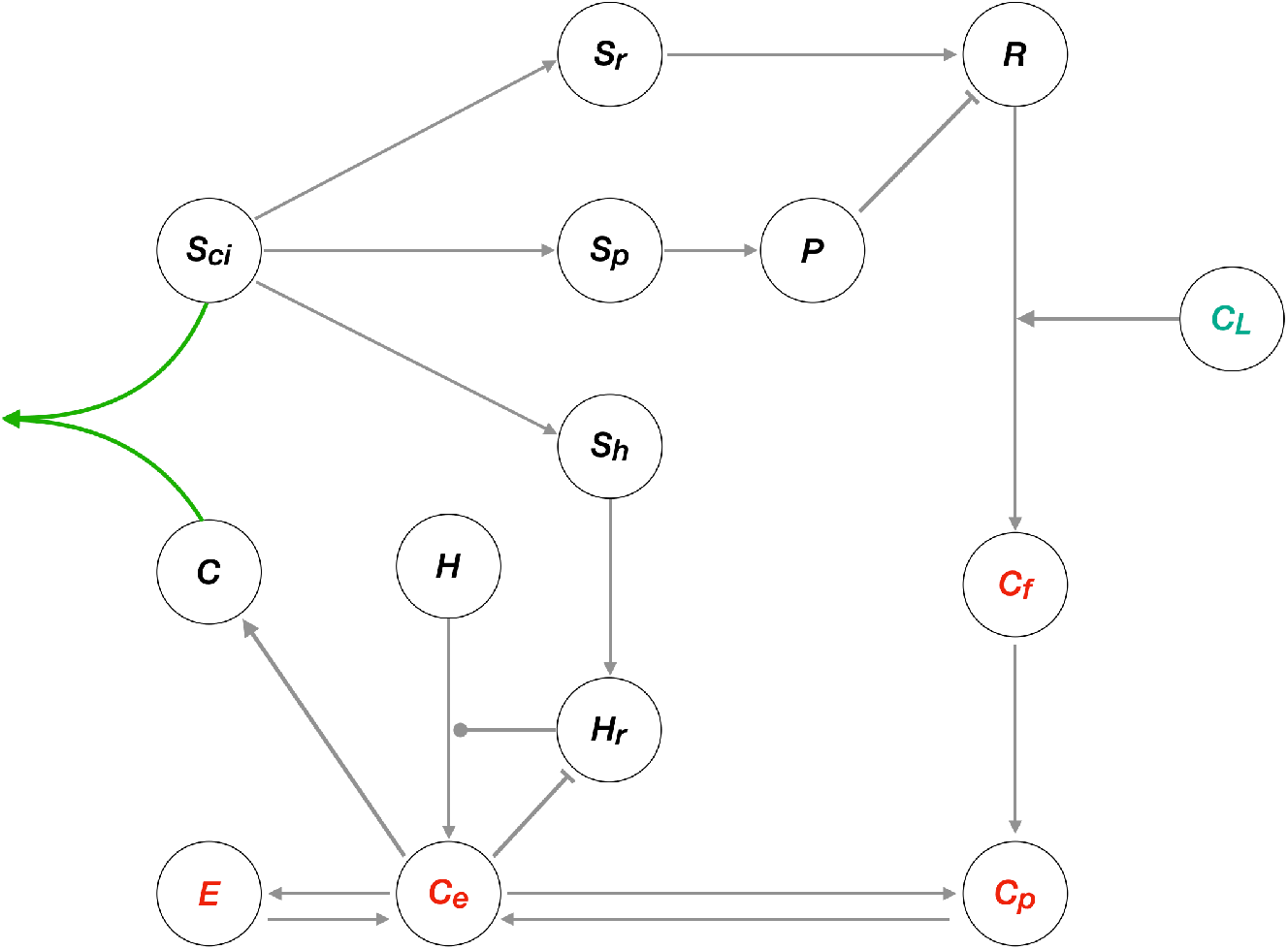
Network schematic for the cellular cholesterol regulatory network. This network diagram captures the nature of the interactions among the thirteen species of the CRN, illustrating the overarching topology of the cellular cholesterol homeostatic machinery as a single Opposer module, with its characteristic feedback architecture. Grey arrows represent flux, or an activating/upregulating influence, while blunt arrow heads (flat bars) represent inhibition or a negative/downregulating influence. Solid circular line endings represent a catalytic reaction. The single opposer node, comprising the sequestration of cholesterol within the SREBP/Scap/Cholesterol complex (*S*_*ci*_*C*) is represented by the green arrows.

Our graphical analysis suggests that it is the concentration of cholesterol in the ER membrane (*C*_*e*_) that is subject to a molecular control mechanism, and thereby acts as a ‘sensor’ molecule (see Subnetwork 1A). Once RPA is conferred to this specific subcellular cholesterol concentration, RPA is then also conferred to the cholesterol concentration at the cellular plasma membrane (*C*_*p*_), to the cholesterol concentration released form the LDLR (*C*_*f*_) and to the esterified cholesterol concentration (*E*), by virtue of the relationships of these molecules to *C*_*e*_ in the CRN graph structure (see Subnetwork 1A). Molecular concentrations in the remaining independent subnetwork of the CRN (see Subnetwork 2) are not subject to RPA.

By computing a projection of the ideal generated by the mass-action equations onto two variables — the input to the network, *C*_*L*_, and the RPA-capable sensor molecule, *C*_*e*_ — we demonstrate that the simple linear coordinate transformation that extracts an RPA polynomial from the system is unique, and corresponds to a known and well–characterised molecular control mechanism known as antithetic integral control (Briat et al., 2016; Cappelletti et al., 2020). This simple RPA-conferring control mechanism has previously been identified in the form of sigma/anti-sigma factors in bacteria, and has also been incorporated in synthetic implementations of homeostatic control (Aoki et al., 2019). Our study now reveals that antithetic integral control is almost certainly the basis for the exquisitely controlled concentration of cholesterol in cellular and sub-cellular membranes. In this case, the counterpart to the interaction between sigma and anti–sigma factors is the sequestration of active cholesterol (*C*) by the SREBP/Scap/Insig complex (*S*_*ci*_). The linear combination of reaction rates involving these two variables gives rise to the single opposer node of the Opposer module, which we highlight in greater detail in Figure 6. Our analysis is also able to explicitly determine all setpoints for RPA-capable molecules as functions of network parameters.

**Figure 6.**
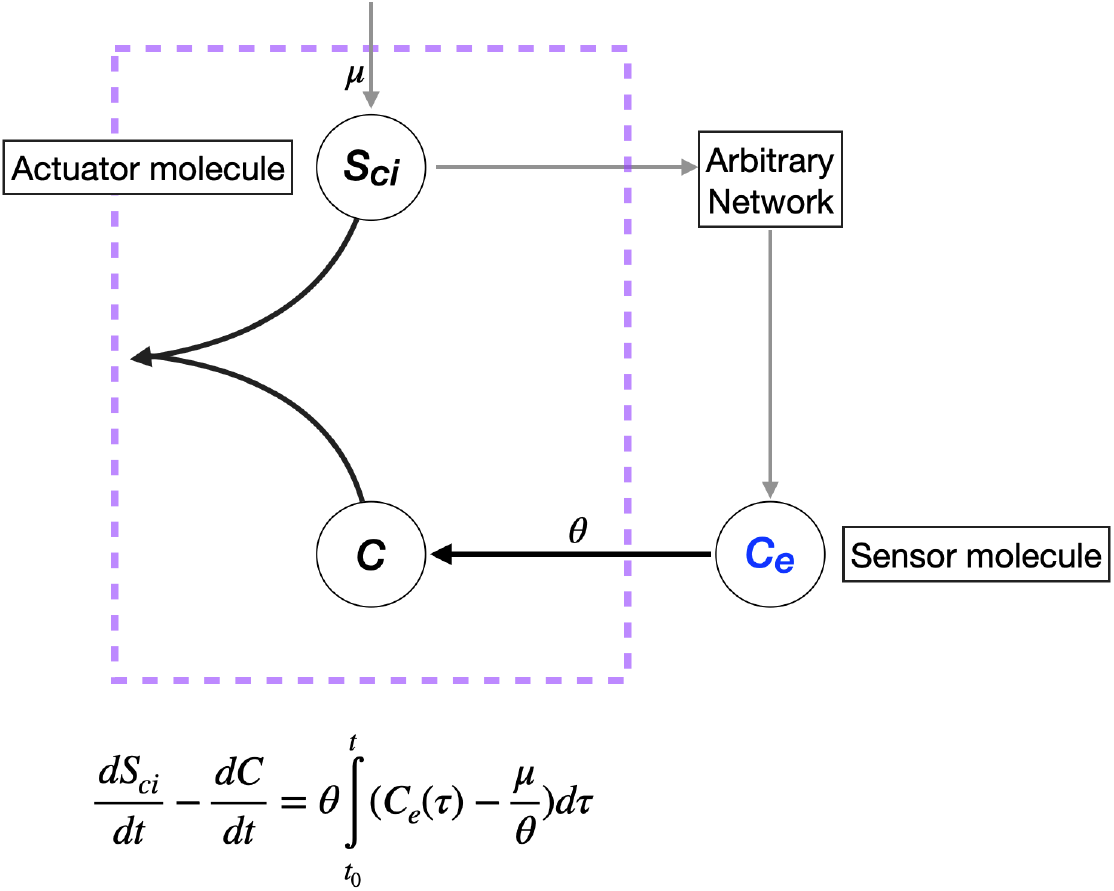
Single opposer node for the cellular cholesterol regulatory network. The purple box highlights the sequestration process that implements a form of antithetic integral control, thereby constituting a single opposer node via a single linear coordinate change. These opposer interactions confers RPA on the sensor molecule, *C*_*e*_, which in turn imparts RPA to several upstream molecules (not shown).

The modelling and analysis approach presented here demonstrates how the fundamental network design principles required for robust homeostatic control can be realised within a relatively complex cellular regulatory network. Our study provides a detailed framework for exploring the necessary biochemical conditions for robust homeostasis and adaptation in other tightly regulated molecules, or sensory molecules, in complex networks throughout biology.

## AUTHOR CONTRIBUTIONS

R.S.: conceptualization, formal analysis, methodology, software, visualization, writing: original draft preparation. R.P.A.: conceptualization, methodology, writing: review and editing, supervision.

Data Accessibility: No data collected in this study.

Code Accessibility: All code developed and used in this study is available at: https://github.com/RonelScheepers/RobustPerfectAdaptation.

## ACKNOWLEDGEMENTS

R.P.A. is supported by an Australian Research Council (ARC) Future Fellowship (project no. FT190100645) from the Australian Government.

