## Supplementary Material for "Robust Homeostasis of Cellular Cholesterol via Antithetic Integral Control"

### Supplementary information for Robust Homeostasis of Cellular Cholesterol via Antithetic Integral Control (R. Scheepers and R. P. Araujo)

#### 1 Accessing the open–source software Singular

The open–source software Singular can either be accessed via Jupyterlab, an extensible environment for interactive and reproducible computing that is based on the Jupyter Notebook and architecture, or via running Singular from the computer's Terminal. In the first instance, follow the steps below to run Singular as a web–based interface on a local computer:

1. Download Anaconda from https://www.anaconda.com/products/distribution,
2. Open Anaconda Navigator from applications or through search,
3. Open JupyterLab from Navigator,
4. Install Jupyter-kernel-singular on terminal by copying code on https://anaconda.org/conda-forge/jupyter-kernel-singular,
5. In the launcher, select Singular under the Notebook tab. The active file is now ready for the code to be entered.
6. If the code does not run, install Python by following the instructions on the Singular homepage http://www.singular.uni-kl.de/, under the Graphical Interface tab,
7. Finally, check that Python is in PATH by opening Terminal and typing the command: echo $PATH.

To run Singular from the Terminal, the application can be downloaded from https://www.singular.uni-kl.de/index.php/singular-download.html

[Decker et al.(2022)Decker, Greuel, Pfister, and Schönemann]. The results found in this study can be replicated by running the code listings provided in the .txt files on Github into the Singular interface on Terminal.

### 2 Singular code for analysing the RPA capacity of the cholesterol chemical reaction network

#### 2.1 RPA setpoint of $C_e$

By implementing the *Singular* script developed and explained in the supplementary material accompanying [Araujo and Liotta(2023)], we can systematically test the RPA capacity of any variable in the cellular cholesterol network as follows:

1. When analysing CRNs, the relevant underlying field is the real numbers, with characteristic zero. We define a ring, F, where the '0' noted before the parameter list denotes this characteristic. The parameters are listed in any order, while the variables list has the two projection variables we want to test ($C_e$ and $C_L$), at the end. The command '(dp(11),dp(2))' denotes the monomial block elimination order (dp($n-2$), dp(2)).This elimination method imposes the computationally efficient *degree reverse lexicographic order* on both the block of $(n-2)$ variables to be eliminated and the block of 2 variables that remain (See Supplementary Material in [Araujo and Liotta(2023)]). The code is entered as:

```
ring F=(0,k1,k2,k3,k4,k5,k6,k7,k8,k9,k10,k11,k12,k13,k14,k15,k16,p1,p2,p3,
mu,eta,theta,alpha),(Sp,C,P,R,Sr,Sh,H,HR,Cp,E,Sci,Cf,Ce,CL),(dp(11),dp(2));
```

2. Next, the mass–action rate equations is listed as:

```
poly f1 = mu - eta*Sci*C;
poly f2 = theta*Ce - eta*Sci*C;
poly f3 = k1*Sci - k2*Sr;
poly f4 = k1*Sci - k10*Sh;
poly f5 = p1*Sr - k11*R - k14*P*R;
poly f6 = k3*CL*R - k4*Cf;
poly f7 = k4*Cf - k5*Cp + k6*Ce;
poly f8 = k5*Cp - k6*Ce + k9*HR*H - k12*Ce - k7*Ce + k8*E;
poly f9 = k7*Ce - k8*E;
poly f10 = p2*Sh - k15*HR*Ce;
poly f11 = alpha - k9*HR*H;
poly f12 = k1*Sci - k13*Sp;
poly f13 = p3*Sp - k16*P;
```

3. To compute the elimination ideal generated by the rate equations $f_1, ...., f_{13}$, we begin with the command:

```
ideal I = f1, f2, f3, f4, f5, f6, f7, f8, f9, f10, f11, f12, f13;
```

   Next, the Gröbner basis for the ideal with the monomial ordering selected in step (1) is computed via:

```
ideal GI = groebner(I);
GI;
```

The CRN is RPA-capable if and only if the first polynomial in the Gröbner basis (listed as GI[1]) is the only generator that comprises only the two projection variables, and it must be factorizable into an RPA polynomial, $\rho$.

4. Finally, a set of polynomials $\{r_1, ...r_n\}$ such that $r_1 f_1 + r_2 f_2 + ...r_n f_n = \rho =$ GI[1] is calculated via the *lift* command to reveal which linear combination of the original rate equations 'reaches' the RPA polynomial:

```
lift(I,GI[1]);
```

In this case, the RPA polynomial $\rho = GI[1] = -f_1 + f_2$ since $r_1 = -1$ and $r_2 = 1$ ($r_3 = r_4 = ... = r_{13} = 0$). This gives $-f_1 + f_2 = \theta C_e - \mu$. The setpoint is then computed as:

$$C_e = \frac{\mu}{\theta}$$

5. To determine if any conservation laws exist, the syzygies of our system are computed through the command:

```
syz(I);
```

While the output is not shown here due to extensive amount of code generated, it can be concluded that no conservation laws exist for the cholesterol system as no linear synergies (generators containing only constant elements) have been generated. RPA for this system is thus uniquely conferred by the linear change of coordinates that produces the specific RPA polynomial.

Similarly, we can now test the RPA capacity of any other variable in the cellular cholesterol network by projecting it against the known non–RPA variable, the input to the system $C_L$. The full code listings to determine the set point values of the RPA–variables $C_e, C_p, C_f$ and $E$ are available on https://github.com/RonelScheepers/RobustPerfectAdaptation.
